# Evaluation of non-modified wireframe DNA origami for acute toxicity and biodistribution in mice

**DOI:** 10.1101/2023.02.25.530026

**Authors:** Eike-Christian Wamhoff, Grant A. Knappe, Aurora A. Burds, Rebecca R. Du, Barry W. Neun, Simone Difilippantonio, Chelsea Sanders, Elijah F. Edmondson, Jennifer L. Matta, Marina A. Dobrovolskaia, Mark Bathe

## Abstract

Wireframe DNA origami can be used to fabricate virus-like particles for a range of biomedical applications, including the delivery of nucleic acid therapeutics. However, the acute toxicity and biodistribution of these wireframe nucleic acid nanoparticles (NANPs) have not previously been characterized in animal models. In the present study, we observed no indications of toxicity in BALB/c mice following therapeutically relevant dosage of unmodified DNA-based NANPs via intravenous administration, based on liver and kidney histology, liver biochemistry, and body weight. Further, the immunotoxicity of these NANPs was minimal, as indicated by blood cell counts and type-I interferon and pro-inflammatory cytokines. In an SJL/J model of autoimmunity, we observed no indications of NANP-mediated DNA-specific antibody response or immune-mediated kidney pathology following the intraperitoneal administration of NANPs. Finally, biodistribution studies revealed that these NANPs accumulate in the liver within one hour, concomitant with substantial renal clearance. Our observations support the continued development of wireframe DNA-based NANPs as next-generation nucleic acid therapeutic delivery platforms.

## Introduction

DNA nanostructures have been extensively explored in biomedical applications^1-3^, and have emerged as a promising delivery platform for vaccines^4-6^, nucleic acid therapeutics^7-10^, and small molecule drugs^11-13^. The DNA origami method can produce monodisperse DNA-based nucleic acid nanoparticles (NANPs) on the 10-100 nm length scale with near-quantitative yields^14^. Compared with other nanoparticle (NP) delivery platforms, NANPs have several unique advantages^3, 15^. Independent control over the size and geometry of NANPs with site-specific functionalization may improve tissue- and cell-specific targeting^16^, and their sequence-based programmability can be leveraged for logic-gated, controlled cargo release^17, 18^. NANPs have tunable biodegradation profiles^19^, and can be modified chemically and structurally to control immunostimulation^4, 20, 21^. Additionally, NANPs can simply incorporate combinatorial ratios of nucleic acid therapeutics including siRNAs and CRISPR-RNPs through nucleic acid hybridization^15^.

Two major classes of DNA origami have been developed to date: dense brick-like^22-24^ and porous wireframe^25-27^ architectures. Comparatively, wireframe NANPs require less nucleic acid for fabrication of similar size objects, which might lead to less immunostimulation, and additionally are stable in physiological buffers^19, 28^. Employing top-down design algorithms, wireframe NANPs can be rapidly prototyped in both two^29-33^ and three dimensions^28, 32-35^. Fully scalable production methods of single-stranded DNA ‘scaffolds’ with programmable sequence and length now enable preclinical and clinical biomedical applications for both classes of DNA origami^36-40^.

DNA-based NANPs are biodegradable via endogenous nucleases, most notably DNase I extracellularly^19, 41^. This limits their potential cardio and pulmonary toxicity, which is associated with other nanoparticles^42^, although this characteristic can also limit *in vivo* circulation times^19, 41,43^. Two of the three major toxicity mechanisms of NPs, namely reactive oxygen species generation and release of metal ions, are not relevant for NANPs. Immunotoxicity has emerged as the third major toxicity type. Notably, both single- and double-stranded DNA (ssDNA and dsDNA) are recognized by the immune system through pattern recognition receptors. ssDNA containing unmethylated cytosine-guanine dinucleotide (CpG) sequence motifs activates endosomal toll-like receptor 9 (TLR9) signaling, and dsDNA activates the cytosolic cyclic GMP-AMP synthase (cGAS)-stimulator of interferon genes (STING) pathway^44, 45^. Additionally, NLRP3, AIM2, and IFI16 are components of the inflammasome capable of recognizing sxs- and dsDNA^45^. The induction of proinflammatory cytokines via these signaling pathways can potentially result in hypersensitivity reactions like cytokine storm and anaphylaxis^46, 47^. Furthermore, antibody responses against nuclear DNA and chromatin have been implicated in driving autoimmunity and the pathogenesis of systemic lupus erythematosus (SLE)^48^. Thus, the potential for immunostimulation and immunotoxicity is a major safety consideration for DNA origami in biomedical applications.

Yet, the immunotoxicity of DNA-based NANPs remains underexplored, particularly in animal models and for wireframe NANPs. Initial *in vitro* cell-based studies revealed context-dependent immunostimulation with examples of rod-like DNA origami being immunologically inert^49^, contrasted by modest immune cell activation by rod-like^20^ and wireframe DNA origami^21,50^. Both TLR9 and the cGAS-STING pathways contributed to immune recognition^20, 21^. Additionally, immunostimulation can intentionally be enhanced by the multivalent display of specific immunostimulatory nucleic acids, both *in vitro*^21, 49^ and *in vivo*^4^. To date, however, no indications of immunotoxicity have been observed in animal models, with studies exploring different NANP geometries and administration routes, namely intravenous (i.v.)^12, 18, 51, 52^ and intraperitoneal (i.p.)^52^. NANPs dosed below 1 mg/kg were reported to be immunologically inert^12, 18, 51^ while more therapeutically relevant doses induced modest immunostimulation^52, 53^ (for reference, therapeutic nucleic acids are typically dosed at 1-10 mg/kg clinically^54^). Several studies additionally evaluated the NANP-mediated induction of total and DNA-specific antibody responses^6, 52, 55^ (i.e., immunogenicity). While total IgM levels transiently increased, no indications of DNA-specific immunological memory or autoimmunity were observed. In contrast to brick-like NANPs, the immunotoxicity and immunogenicity of wireframe NANPs have not been characterized at therapeutically relevant doses in animal models.

In addition to preclinical safety profiles, characterizing the biodistribution is essential for biomedical applications. Several brick-like DNA origami biodistribution studies have been conducted^10, 12, 18, 50-52^, exploring various shapes, routes of administration, and both healthy as well as tumor-bearing mouse models. Again, the biodistribution of wireframe DNA origami remains underexplored. One study investigated the biodistribution of a wireframe NANP with six helices per edge after i.v. administration in C57 nude mice, and observed rapid bladder accumulation and an elimination half-life under one hour^50^.

Here, we characterize the acute toxicity and biodistribution of wireframe DNA-based NANPs following i.v. and i.p. administration. In BALB/c mice at 4 mg/kg doses, we found no phenotype across animal body weight, liver and kidney histology, and liver biochemistry. Additionally, we observed no phenotype in immune blood cell counts, and minimal cytokine induction. In SJL/J mice, commonly used as a model of autoimmunity, i.p. administration of NANPs did not induce DNA-specific IgM or IgG antibody production; consequently, immune complex-mediated kidney damage was also not observed. Finally, upon i.v. administration of these NANPs, we observed accumulation in the liver within one hour, accompanied by rapid renal clearance, as anticipated for biodegradable nanomaterials^56, 57^. Taken together, our study suggests that wireframe DNA origami have limited immunotoxicity and accumulate in the liver, supporting their continued development for biomedical applications including both liver-targeting of nucleic acid therapeutics and also addressing the grand challenge of engineering these materials for extrahepatic delivery.

## Results

We designed and conducted a set of animal model experiments to investigate the biodistribution, acute toxicity, and, specifically, immunotoxicity of wireframe NANPs (**Figure 1.A**). We administered 4 mg/kg NANP i.v. into BALB/c mice and investigated toxicity readouts including body weights, liver and kidney histology, liver and kidney biochemistry, blood cell counts, and cytokine induction. Additionally, we administered 0.4 mg/kg NANP into SJL/J female mice i.p. to understand the potential for wireframe NANPs to break immunological tolerance to self-DNA and induce an autoimmune response. Finally, we administered 2 mg/kg of fluorophore-labeled NANP into BALB/c mice i.v. to characterize their biodistribution. Taken together, these experiments provided a baseline characterization of wireframe NANP toxicity and biodistribution at therapeutically relevant doses.

**Figure 1.**
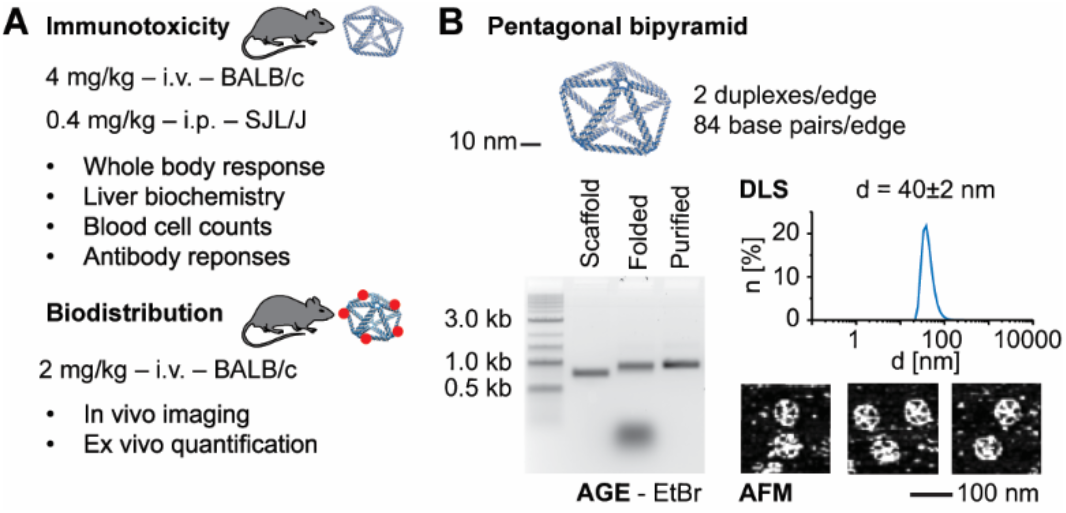
Characterizing the acute toxicity and biodistribution of wireframe DNA origami. **(A)** The experimental design for this study investigated the acute toxicity, immunotoxicity and biodistribution of wireframe NANPs across two different animal models via two different administration routes. **(B)** The model NANP, a pentagonal bipyramid with two duplexes and 84 base pairs per edge (**PB84**), used in this study. **PB84** structural integrity was validated by agarose gel electrophoresis (AGE), dynamic light scattering (DLS), and atomic force microscopy (AFM).

We designed and fabricated a pentagonal bipyramid with 84 base pairs and two DNA duplexes per edge (**PB84**) to serve as a model to assess the immunotoxicity and biodistribution of wireframe NANPs (**Figure 1.B** and **S1**). Characterizing **PB84** by agarose gel electrophoresis, dynamic light scattering, and atomic force microscopy, we observed monodisperse nanostructures with an approximate diameter of 40 nm, which we were able to prepare in milligram quantities for subsequent *in vivo* experiments.

We first administered 4 mg/kg of **PB84** i.v. in at least four female BALB/c mice for each monitored toxicity readout. Monitoring animal body weights over seven days after administration, no weight loss was observed, comparable to PBS and unstructured ssDNA controls (**Figure 2.A**). Next, we examined liver and kidney histology, noting that these two organs were the major sites of accumulation observed in previous DNA origami studies^12, 18, 50-52^. We observed no indications of pathology in the liver and kidney (**Figure 2.B**). We assayed liver and kidney biochemistry after administration of **PB84** and observed no phenotype for the following biomarkers for toxicity: alkaline phosphatase, aspartate aminotransferase, alanine aminotransferase, blood urea nitrogen, creatinine, total serum protein, albumin, globulin, glucose, and bilirubin (**Figure 2.C**).

**Figure 2.**
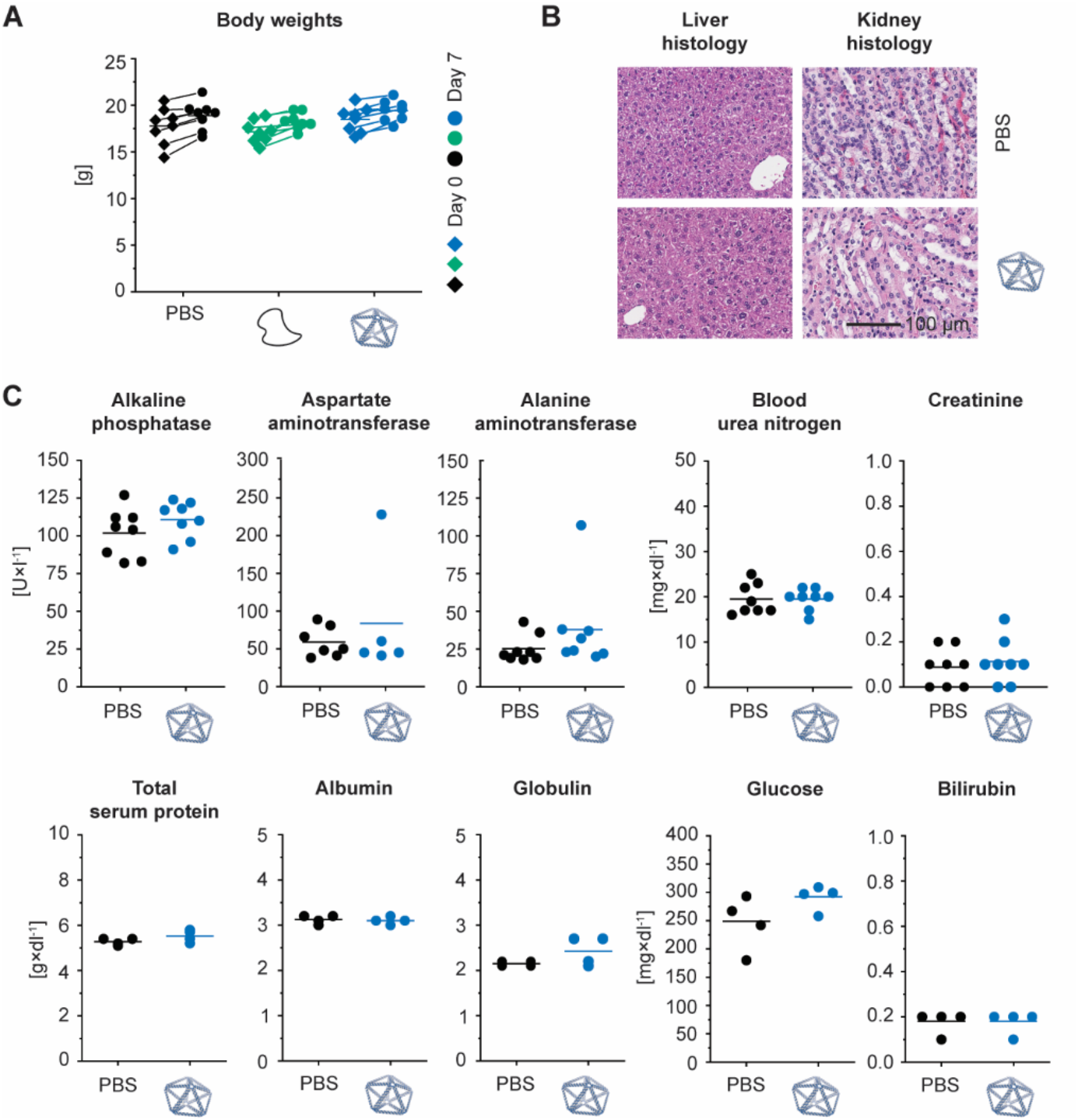
Effect of wireframe DNA origami on body weight as well as liver and kidney histology and function in BALB/c mice. Intravenous administration of PB84 (4mg/kg), PBS control and an unstructured DNA control (4mg/kg). **(A)** Body weights were monitored before and seven days after administration. Body weight change was consistent across all groups. **(B)** Histological sections of kidney and liver were visualized by hematoxylin and eosin (H&E) staining and observed under a light microscope at 40× magnification. **(C)** A panel of 10 biomarkers for liver and kidney function shows no phenotype when NANPs were administered, consistent with a PBS control. General toxicity was assessed from n ≥ 4 biological replicates per group. Representative histology images are shown. Student’s t-test was performed for the body weights. One-way ANOVA was performed for the liver and kidney biochemistry panel followed by Tukey’s multiple comparison test. Significant differences are denoted as * - p < 0.050, ** - p < 0.010, and *** - p < 0.001.

Next, we investigated whether NANP administration resulted in the proliferation of blood cell types. No phenotype for abnormal cell counts was observed for total white blood cells, as well as for lymphocytes and monocytes individually (**Figure 3** and **S2**). In addition to blood cell counts, we investigated whether cytokines were produced following **PB84** administration, as they have been implicated in NP and nucleic acid immunotoxicity^42, 47^. We conducted an enzyme-linked immunoassay (ELISA) with serum from blood drawn at 3 and 24 hours after administration, reflecting differential kinetics of cytokine induction (**Table 1**). We observed IL-6 and CXCL2 induction at 3 hours, and IL-12 and TNF induction at 24 hours, although these cytokine levels were not significantly increased compared to the PBS control.

**Table 1.**
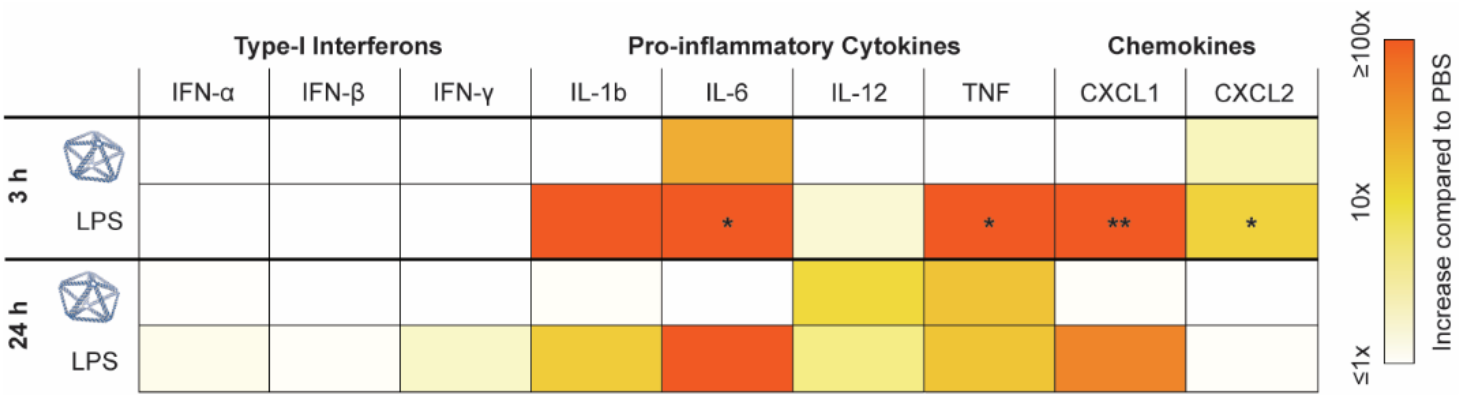
Cytokine levels in BALB/c mice after i.v. administration. A cytokine array panel was conducted at 3 and 24 hours after administration. PBS, **PB84**, and lipopolysaccharide (LPS) were administrated i.v., and serum was collected at corresponding time points. Cytokine induction was assessed from n = 3 biological replicates per group. Cytokine induction compared to PBS is shown. One-way ANOVA was performed for the 3 h and 24 h time point followed by Tukey’s multiple comparison test. Significant differences compared to PBS are denoted as * -p < 0.050, ** -p < 0.010, and *** -p < 0.001.

**Figure 3.**
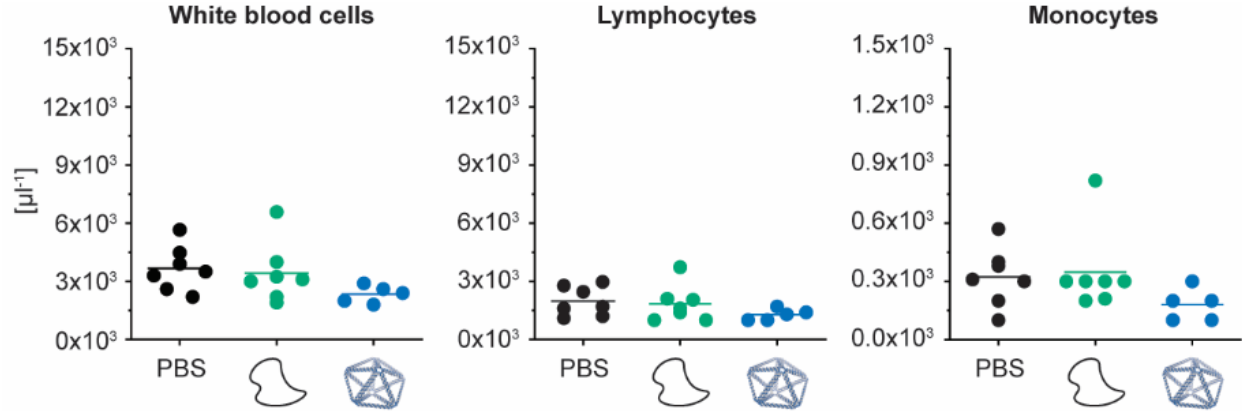
Characterizing blood cell counts in BALB/c mice after i.v. administration. BALB/c mice were intravenously administrated through tail-vein injection with 4 mg/kg of **PB84** per animal. Total white blood cell, lymphocyte, and monocyte cell counts were not elevated when NANPs were administered, consistent with a PBS control and an unstructured ssDNA control. Blood cell counts were assessed from n ≥ 5 biological replicates per group. One-way ANOVA was performed for the blood cell counts followed by Tukey’s multiple comparison test. Significant differences are denoted as * - p < 0.050, ** - p < 0.010, and *** - p < 0.001.

Next, to evaluate the potential of NANPs to break immunological tolerance to self-antigens and induce an autoimmune response, we investigated their safety profiles in the SJL/J mouse model. This mouse model is genetically predisposed to develop autoimmunity and is commonly used to study chemically induced autoimmunity^58^. Female SJL/J mice have greater mortality and show greater blood levels of DNA-specific antibodies and, subsequently, immune complex-mediated kidney damage than male mice. This phenomenon is similar to SLE in humans, in which the male-to-female disease incidence ratio is 1-to-9, and the disease is more severe in females. Therefore, female mice were used in our study. We injected 0.4 mg/kg of **PB84** i.p. and observed no significant phenotype when assessing kidney histology via a kidney score when compared with a PBS control (**Figure 4.A** and **S3**). In contrast, all animals treated with pristane, used as a positive control, showed histological changes characteristic of immune complex-mediated kidney damage. We additionally monitored the production of DNA-specific IgM and IgG antibodies over 20 weeks (**Figure 4.B**). Unlike pristane, **PB84** did not induce DNA-specific IgM and IgG production.

**Figure 4.**
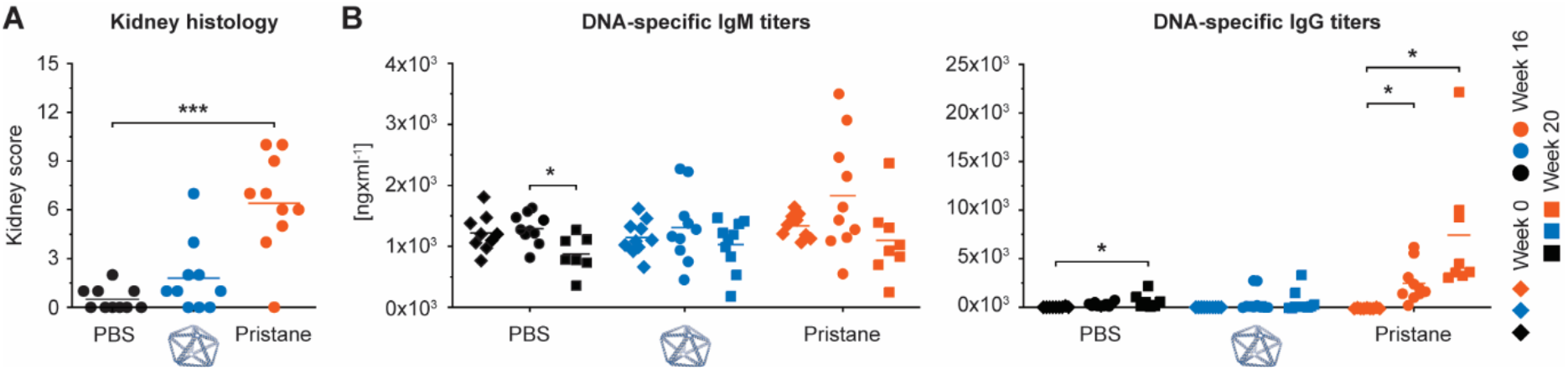
Characterizing autoimmunity induction in SJL/J mice after i.p. administration. SJL/J mice were intraperitoneally administered with 0.4 mg/kg of **PB84** per animal. **(A)** Kidney scores were assessed at 20 weeks post treatment and **(B)** DNA-specific IgM and IgG antibody titers were assessed before treatment (time 0 or baseline) and after 16- and 20-weeks post treatment from ten animals (n = 10) per group. One-way ANOVA was performed followed by Tukey’s multiple comparison test. Significant differences are denoted as * - p < 0.050, ** - p < 0.010, and *** - p < 0.001.

Given that **PB84** was generally safe in different animal models and via different modes of administration, we sought to further understand this nanomaterial’s potential as a delivery vehicle. Towards this end, we conducted a baseline biodistribution study of **PB84**. To facilitate *in vivo* and *ex vivo* characterization of **PB84**’s biodistribution, we installed 5 Alexa Fluor 750 fluorophores via pre-assembly functionalization to yield **PB84-5xAF750**. Following i.v. administration at 2 mg/kg, **PB84-5xAF750**’s biodistribution was monitored for several hours (**Figure 5.A** and **S4**) by *in vivo* imaging and organ accumulation was characterized *ex vivo* after 60 minutes (**Figure 5.B** and **S5**). Within 15 minutes, **PB84-5xAF750** was rapidly cleared from circulation as measured by blood draws (**Figure 5.C**), where it mainly accumulated in the liver (**Figure 5.A** and **S4.B**) and remained for at least two hours as renal clearance initiated. Importantly, this biodistribution profile was different than a fluorophore-labelled, control oligonucleotide, which was renally cleared within 15 minutes, with substantially less accumulation in the liver. *Ex vivo* imaging of harvested organs confirmed that **PB84-5xAF750** mainly accumulates in the liver after 60 minutes, with minor accumulation in the spleen, kidney, and lung (**Figure 5.B** and **S5**).

**Figure 5.**
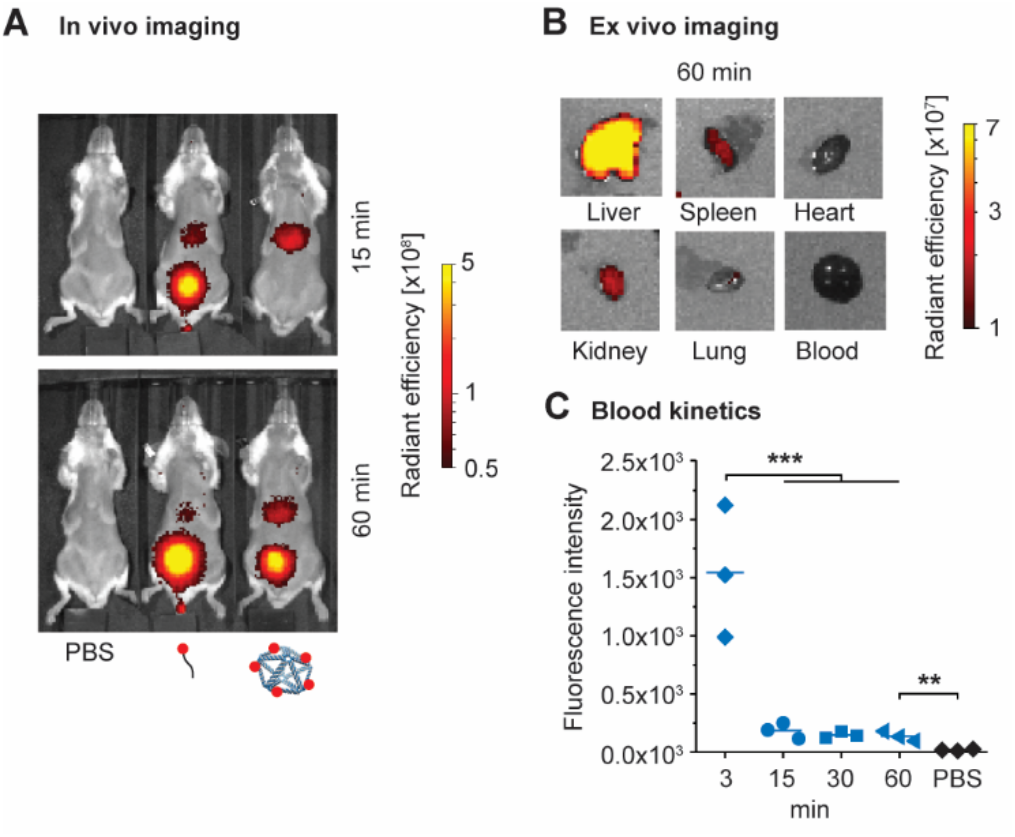
Biodistribution in BALB/c mice after i.v. administration. In vivo optical imaging for analysis of pharmacokinetics and biodistribution. BALB/c mice were injected with PBS, AlexaFluor750-labeled oligonucleotide, or a DNA origami **PB84-5xAF750** and imaged at multiple timepoints post-injection. **(A)** Representative fluorescence images at 15- and 60-minutes post-injection (n=3 per group). **(B)** Representative fluorescence images of ex vivo organs harvested 60 minutes post-injection of **PB84-5xAF750. PB84-5xAF750** was mostly accumulated in the liver, with minor accumulation in the kidney, spleen, and lung after 60 minutes. **(C)** Fluorescence intensity measurements of drawn blood for up to 60-minutes post-injection. One-way ANOVA was performed for blood kinetics measurements followed by Tukey’s multiple comparison test. Significant differences are denoted as * - p < 0.050, ** - p < 0.010, and *** - p < 0.001.

## Discussion

Overall, these data indicate that unmodified wireframe DNA-based NANPs are not acutely toxic after a single i.v. administration of therapeutically relevant doses, and NANPs accumulate in the liver prior to biodegradation and renal clearance. Unlike earlier studies with antisense DNA oligonucleotides which reported mononuclear cell infiltration in the liver and kidney^59^, our general toxicity study did not reveal any kidney or liver damage as indicated by histology. We did not observe any phenotypes when assessing liver and kidney biochemistry, suggesting that NANPs do not alter the healthy functioning of the liver, even though this is their primary site of accumulation. We observed no change in the absolute globulin levels and albumin-globulin ratios. This indicates there was no onset of hypergammaglobulinemia, a pathology characterized by the over-production of globulin proteins by B cells, which is a characteristic response to antisense DNA oligonucleotides^59^. Monocytosis, another phenotype linked with antisense DNA oligonucleotides^59^, was not observed either. This is in contrast to a previous study^52^ in which transient monocytosis was observed for brick-like NANPs (12 mg/kg, i.p. administration, five repeat doses), which might be due to differential cell-specific uptake by these dense objects compared with the porous wireframe architecture explored here^60^.

Importantly, mouse models generally serve as conservative estimates for immune cell proliferation, such as monocytosis and splenomegaly, in response to nucleic acid therapeutics. Only when tested in animals genetically predisposed to autoimmunity (SJL/J females), NANPs resulted in grade 3 spleen plasmacytosis in one of ten animals (data not shown); since one animal with grade 2 plasmacytosis was also detected in the PBS negative control group (data not shown), this observation suggests that the risk of increase in the number of immune cells in response to NANPs is low.

Since cytokine induction has been implicated in nanomaterial toxicity^42, 47^, this innate immune response was important to characterize. Three hours after administration, we observed IL-6 and CXCL2 induction, and at 24 hours, we observed IL-12 and TNF induction; this induction was not significantly higher than in the PBS control. Similar mild immunostimulation has been observed in other DNA origami studies^52, 53^. DNA-induced TLR-9 signaling can lead to the production of IL-6, IL-12, and TNF^45^. However, while TLRs have often been implicated in nucleic acid therapeutic immunostimulation, they are often not the only source^59^. Further studies are required to mechanistically determine the origins of the observed cytokine production. We note that for some applications, such as cancer immunotherapy, the ability to stimulate proinflammatory cytokine production might be a preferable characteristic of a delivery vehicle. Researchers have demonstrated proof-of-concepts that immune system stimulation can be programmed with NANPs^4, 20, 21, 49^. We anticipate this capability will be important in future biomedical applications.

One potential safety concern relevant to the *in vivo* use of DNA-based NANPs is breaking the immunological tolerance to DNA and inducing autoimmune responses via the generation of DNA-specific antibodies. While no ideal preclinical model for autoimmunity studies is currently available, SJL/J mice are commonly used to model the development of chemically-induced SLE, an autoimmune condition characterized by broad self-reactive antibody responses and glomerulonephritis^58^. We found that, unlike pristane, used as a positive control, NANPs do not induce SLE phenotype in the SJL/J model, as assessed by kidney histology and DNA-specific IgM and IgG serum levels. These data corroborate additional findings that DNA origami generally does not elicit DNA-specific immunological memory^6, 52, 55^. We note that further understanding the adaptive immune response against NANPs at different doses, dosing regimens, and routes of administration will be important to further validate these findings. NANPs co-formulated with peptides and proteins may also alter these adaptive immune responses.

A preliminary biodistribution study revealed that these wireframe NANPs rapidly exit the bloodstream on the order of minutes, mainly accumulating in the liver at 60 minutes post-administration. At 4 hours (**Figure S4.B**), the NANPs were cleared from the liver, suggesting that the NANPs are biodegrading in the liver or bloodstream into fragmented NPs or oligonucleotides that can then be processed by the renal system. This biodistribution is consistent with previous studies on brick-like NANPs that found the liver as a major site of accumulation for NANPs^12, 18, 50, 52^, as well as numerous studies of different classes of NPs due to the drastic reduction in hepatic blood flow velocity and fenestrated cellular environment^56^. In a notable exception, researchers observed that the kidney was the main organ of accumulation for larger (∼100 nm characteristic length scale) NANPs^51^. Additionally, the biodistribution of DNA origami can be altered when NANPs are functionalized with biomacromolecules^43, 50^. Taken together, these data suggest that the biodistribution of wireframe DNA-based NANPs may also be programmed by modulating the size and shape of the NP, as well as the chemical composition and surface display of molecules.

Overall, these data support the continued development of wireframe DNA origami for biomedical applications. Future investigations could explore various additional directions. Following up on this acute toxicity study, evaluating safety profiles when NANPs are administered across multiple administration routes at increased (and repeated) doses will lead to a deeper understanding of the potential therapeutic window of these nanomaterials. For example, understanding the doses at which safety issues emerge between intravenous and subcutaneous administration will help prioritize further development efforts. We found that upon intravenous administration, NANPs accumulate in the liver. Modulating the size, shape, and surface display of molecules on NANPs may lead to accumulation in extra-hepatic tissues, which is of high priority in the biomedical delivery field. Finally, metabolic studies understanding the rate of biodegradation and the fate of the degradation products are required for better understanding the safety profiles of these NANPs, with and without stabilizing modifications that alter their nuclease-driven biodegradation and may also alter cellular uptake and tissue-targeting properties.

## Acknowledgements

E.-C.W., G.A.K., R.R.D., and M.B. were supported by NSF CCF-1564025, NSF DMREF CBET-1729397, NIH R21-EB026008, NIH R01-MH112694, ONR N00014-17-1-2609, ARO ISN W911NF-13-D-0001, and FastGrant AGMT EFF 4/15/20. E.-C.W. was additionally supported by the Feodor Lynen Fellowship of the Alexander von Humboldt Foundation. G.A.K. was additionally supported by the National Science Foundation under a Graduate Research Fellowship 2389237. This work made use of the MRSEC Shared Experimental Facilities at MIT, supported by NSF DMR-1419807. Additional support for this research was provided by a core center grant P30-ES002109 from the National Institute of Environmental Health Sciences, National Institutes of Health. We thank the Koch Institute’s Robert A. Swanson (1969) Biotechnology Center (SBC) for technical support, specifically the Preclinical Imaging & Testing Facility, the Nanotechnology Materials Facility, and the Histology Facility. We also thank the MIT DCM Comparative Pathology Laboratory. A.A.B and the SBC are supported in part by the Koch Institute Support (core) Grant P30-CA14051 from the NCI. This study was funded in part (B.W.N., M.A.D., E.F.E., and J.L.M.) by federal funds from the National Cancer Institute, National Institutes of Health, under contract 75N91019D00024. The content of this publication does not necessarily reflect the views or policies of the Department of Health and Human Services, nor does mention of trade names, commercial products, or organizations imply endorsement by the US Government.

## Methods

### Materials and Equipment

All materials were purchased from Millipore Sigma, unless otherwise stated. Alexa Fluor 750-NHS was purchased from ThermoFisher Scientific. Nuclease-free water and oligonucleotide staples were purchased from IDT. Agarose was purchased from IBI Scientific. Black 96-well plates were purchased from Nunc. Tris-EDTA (TE), tris-acetate-EDTA (TAE) and phosphate-buffer saline (PBS) were purchased from Corning. The DNA ladder (Quick-Load Purple 2-log DNA ladder 0.1-10 kb) was purchased from New England Biolabs. ToxinSensor Gel Clot Endotoxin Assay Kits were purchased from GenScript. VacciGrade LPS was purchased from InvivoGen.

BioRad T100 Thermal Cyclers were used for DNA nanoparticle assembly. Agarose gels were imaged on a Typhoon FLA 7000. Agarose gel images were processed using ImageJ. DNA concentration measurements were made using a NanoDrop 2000 by ThermoFisher Scientific. Atomic force microscopy (AFM) was conducted using a Veeco Multimode 8 with ScanAsyst-Fluid+ tips. AFM images were processed using ImageJ. Dynamic light scattering (DLS) was performed using a Zetasizer Nano ZSP by Malvern Analytical. Purified oligonucleotides were dried *in vacuo* using a SpeedVac SPD300-DDA by ThermoFisher Scientific. Reversed-phase high-performance liquid chromatography (RP-HPLC) was conducted using a BEH C18 column by Waters. Alfalfa-free mouse diet was purchased from LabDiet. Imaging of live mice and harvested organs was performed using a Caliper Spectrum IVIS and images were process using Living Image Software by PerkinElmer.

### Scaffold and oligonucleotide staple synthesis

The custom-length DNA scaffold (phPB84, 2520 nt) for **PB84** was prepared as previously described^36^. Triton X-144 was used to remove residual endotoxins^43^. Endotoxin levels were measured using ToxinSensor Gel Clot Endotoxin Assay Kits.

To install dyes onto oligonucleotide staples for our biodistribution studies, 50 μM staples containing a 5’ amino groups (IDT’s C6 amino modifier) in PBS at pH 7.4 were reacted with 10x excess of Alexa Fluor 750-NHS dissolved in DMSO overnight at room temperature. Subsequently, excess dye was removed using NAP-10 columns and Alexa Fluor 750-modified staples were purified via RP-HPLC (gradient: 90% 0.1% TEAA and 10% acetonitrile to 10% TEAA and 90% acetonitrile over 30 min). Solvents were removed under vacuum and Alexa Fluor 750-modified staples were dissolved in TE for further use.

### DNA nanoparticle design and assembly

**PB84** and **PB84-5xAF750** were designed using DAEDALUS with the five dyes pointing into the interior of the NANP from the base edges (**Tables S1-S3**)^28^. **PB84** and **PB84-5xAF750** were assembled as previously described^28^. Briefly, a folding reaction containing 30 nM scaffold, 225 nM oligonucleotide staples, 1x TAE, 12 mM MgCl_2_ was prepared in nuclease-free water. The reaction mixture was dispensed into 50 μl aliquots and then subjected to the following thermal annealing treatment: 95 °C for 5 min, 80-75 °C at 1 °C per 5 min, 75-30 °C at 1 °C per 15 min, and 30-25 °C at 1 °C per 10 min. The reaction mixture was then purified into PBS using Amicon Ultra centrifugal filters (100 kDa, 2000 G, 3x, 20 minutes) and stored at 4 °C. Purity and dispersity of resulting NANPs were characterized by agarose gel electrophoresis (1.6 wt% agarose, TAE 12 mM MgCl_2_ buffer, EtBr stain, 65V for 150 minutes at 4 °C) as well as dynamic light scattering. Endotoxin levels were measured using ToxinSensor Gel Clot Endotoxin Assay Kits to ensure administration of less than 5 EU/kg.

### Atomic force microscopy

AFM imaging was performed in TAE with 12 mM MgCl_2_ at pH 8.3 using ScanAsyst mode and ScanAsyst-Fluid tips. Following the deposition of 5 μl **PB84-5xAF750** at 5 nM in PBS at pH 7.4 onto freshly cleaved mica, 2 μl NiCl_2_ at 100 mM were added and incubated for 30 s. Subsequently, 80 μl of TAE with 12 mM MgCl_2_ at pH 8.3 were added to the sample and the AFM tip was submerged into 40 μl TAE with 12 mM MgCl_2_ at pH 8.3 before starting the experiment.

### Animals

Genetically inbred *wild type* male and female BALB/cJ (strain 000651) and female SJL/J (strain 000686) mice were purchased from The Jackson Laboratory (Bar Harbor, ME), and were housed and handled in Association for Assessment and Accreditation of Laboratory Animal Care (AAALAC)-accredited facilities with experimental methods as specifically approved by the Institutional Animal Care and Use Committees at MIT and NCI-Frederick, respectively.

### Gross toxicity, histology, and liver and kidney biochemistry panel in BALB/c model

BALB/c mice received a single 100 μl intravenous tail vein injection consisting of PBS, 4mg/kg unstructured phPB84 scaffold in PBS, or 4mg/kg **PB84** in PBS. Animals were monitored daily for one week. Blood was collected via cardiac puncture to obtain whole blood counts within an hour of collection as well as serum for chemistry panel to assay liver and kidney function (IDEXX BioAnalytics, Columbia, MO). Necropsy was performed to look for signs of gross toxicity. For histologic evaluation, formalin-fixed tissues were embedded in paraffin, sectioned at 5 μm, stained with hematoxylin and eosin (H&E), and visually assessed under a microscope by a veterinary pathologist.

### Cytokine array in BALB/c mice

Cardiac blood was collected from BALB/c mice 3 hr and 24 hr after tail vein injection of 100 μL containing PBS, **PB84** (4mg/kg) in PBS, or immunogenic lipopolysaccharide (LPS 0.05mg/mL). Serum was stored at -80°C before shipment for analysis of induction of cytokines via a custom murine Q-Plex array (Quansys Biosciences). The custom array included IFN-α, IFN-β, IFN-γ, IL-1b, IL-6, IL-12, TNF, CXCL1 and CXCL2.

### Biodistribution studies in BALB/c mice

Prior to biodistribution studies, mice were kept on an alfalfa free diet (LabDiet, AIN-93M, cat# 58M1, irradiated) for 7 days to reduce background fluorescence. BALB/c mice received a single 100ul intravenous tail vein injection consisting of PBS, 2mg/kg oligonucleotide staple-AF750 in PBS, or 2mg/kg DNA origami PB84-5xAF750 in PBS. Live mice were anesthetized with 2% isoflurane and imaged immediately after injection and 15 min, 30 min, 45 min, 1 hr, 2 hr, 4 hr, and 6 hr post-injection. Peripheral blood was collected immediately after injection and 15 min, 30 min, 45 min, and 60 min post-injection, then transferred to a 96-well plate for measurement of the fluorescent signal. For ex vivo imaging of liver, kidney, heart, spleen, and lungs, mice were euthanized via CO_2_ 1 hr post-injection and organs were harvested.

### Autoimmunity study in SJL/J model

Eight weeks old SJL/J females were injected intraperitoneally with 500 μL of either negative control (PBS), positive control (pristane at stock concentration as provided by Sigma Aldrich, catalog # P9622) or 0.4 mg/kg **PB84**. Pristane was synthetic, certified for in vivo studies, having > 95% purity by GC and undetectable endotoxin (< 5EU/mL). Average mouse weight at the time of injection was 19 g. Each treatment group included 10 animals. The blood was collected before injection (baseline) and at 8, 16, and 20 weeks post-injection. Sera were analyzed by ELISA for the presence of anti-dsDNA IgG and IgM using commercially available kits (Chondrex Inc., catalog #3031 and #3032, respectively). At the termination of the study, kidneys were collected, fixed in 4% neutral buffered formalin and stained with Periodic Acid–Schiff (PAS) reagent for detection of immune complex depositions and with hematoxylin and eosin (H&E) for characterization of histologic lesions. Histopathology grading was based on the most severe kidney section for each animal. Grading was performed for glomerular changes, inflammatory infiltrates, and tubular changes as follows: 0 – normal; 1 – minimal; 2 – mild; 3 – moderate; 4 – marked. Inflammatory infiltrates were graded as 0, within normal limits; 1, minimally increased inflammatory infiltrates composed predominantly of lymphocytes and plasma cells often focally forming 3-5 cell thick perivascular cuffs; 2, mildly increased inflammatory infiltrates that are multifocal; 3, moderately increased inflammatory infiltrates that form thick perivascular cuffs multifocally that are prominent with the 4x objective; 4, markedly increased inflammatory infiltrates that widely separate vessels from adjacent renal parenchyma. Tubular changes were graded as follows: 0, within normal limits; 1, minimal tubular degeneration often focal; 2, mild tubular degeneration present with multifocal regions containing tubular degeneration and regeneration and focal necrosis; 3, moderate tubular changes contain tubular degeneration and multifocal necrosis; 4, marked tubular changes including tubular necrosis multifocally. Mesangial expansion was evaluated and graded into four categories: 0, no mesangial expansion; 1, minimal changes; 2, mild mesangial expansion (mesangial matrix wide < 2 nucleus diameter); 3, moderate mesangial expansion often with crescentic glomeruli, mesangial matrix wide < 4 nucleus diameter), and 4, severe mesangial expansion (> 4 nucleus diameter). Cumulative score was calculated and reported as “kidney score”.

## Supplemental Information

Additional AFM data; Additional blood cell count data; SJL/J model histology images; Additional in vivo imaging; Additional ex vivo imaging; Scaffold and staple sequences; Raw data for cytokine panel.

