## Supplementary material for "Evaluation of non-modified wireframe DNA origami for acute toxicity and biodistribution in mice": Supplamental Information

### Supplemental Figures

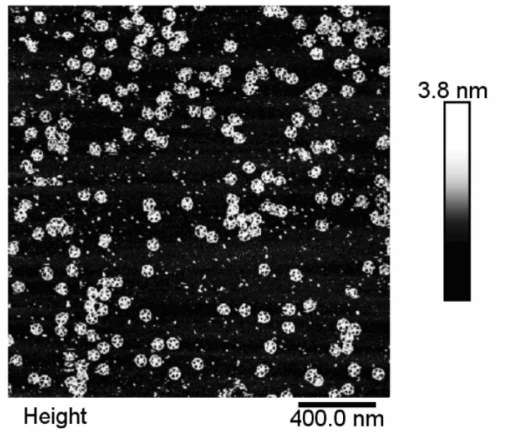


**Figure S1. Supplemental information for atomic force microscopy.**

Representative wide-field atomic force microscopy micrograph of **PB84.**

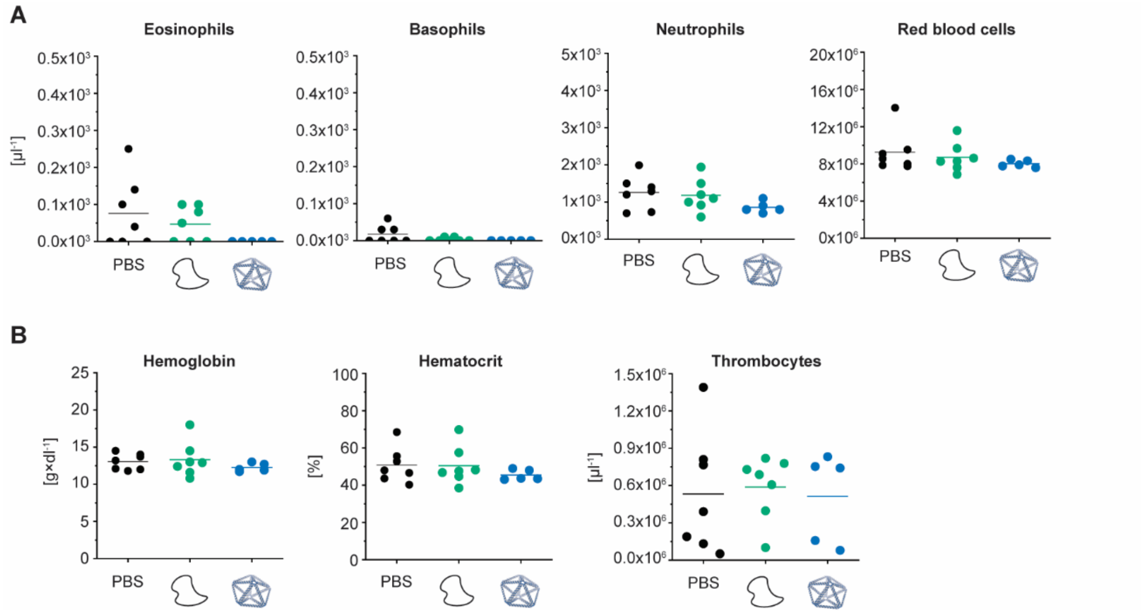


**Figure S2. Supplemental information for blood cell counts in BALB/c mice after i.v. administration.**

BALB/c mice were intravenously administrated through tail-vein injection with 4 mg/kg of **PB84** per animal. **(A)** Eosinophil, basophil, neutrophil, and red blood cell counts were not elevated when NANPs were administered, consistent with a PBS control and an unstructured ssDNA control. **(B)** Hemoglobin, hematocrit, and thrombocytes were not elevated when NANPs were administered, consistent with a PBS control and an unstructured ssDNA control. Blood cell analyses were assessed from $n \geq 5$ biological replicates per group. One-way ANOVA was performed for the blood cell counts followed by Tukey's multiple comparison test. Significant differences are denoted as * - $p < 0.050$, ** - $p < 0.010$, and *** - $p < 0.001$.

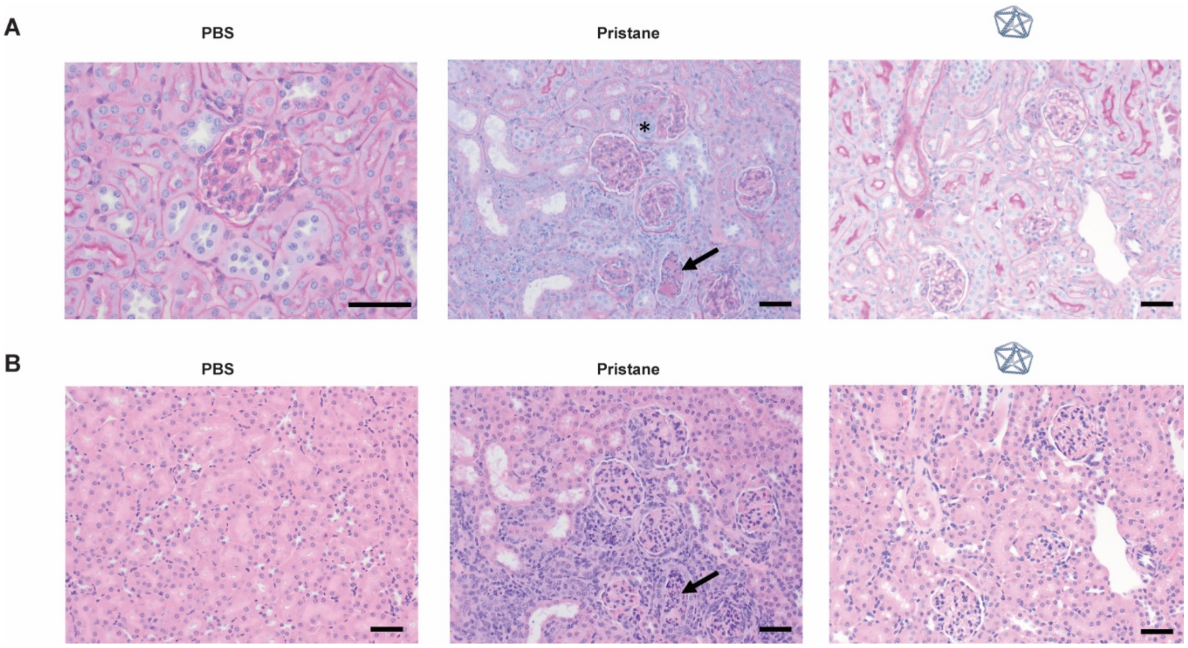


**Figure S3. Supplemental information for autoimmunity induction in SJL/J mice after i.p. administration.**

Representative kidney histology images for the SJL/J mice study. **(A)** PAS stain histology images after PBS, pristane, and **PB84** administration. In the pristane group, glomerular profiles are surrounded by proliferative mesenchymal cells (star) and have increased PAS positive matrix (arrow). Normal glomeruli are within normal limits in the PBS group, and glomeruli are minimally affected in the **PB84** group. **(B)** H&E stain histology images after PBS, pristane, or **PB84** administration. In the pristane group, significant tubular damage is present with degeneration and necrosis of tubular epithelium with dilated tubules containing cellular debris (arrow). Normal tubules of the renal cortex are present in the PBS group, and glomeruli have minimal changes, including minimal glomerular hypercellularity, in the **PB84** group. Scale bars represent 50 µm.

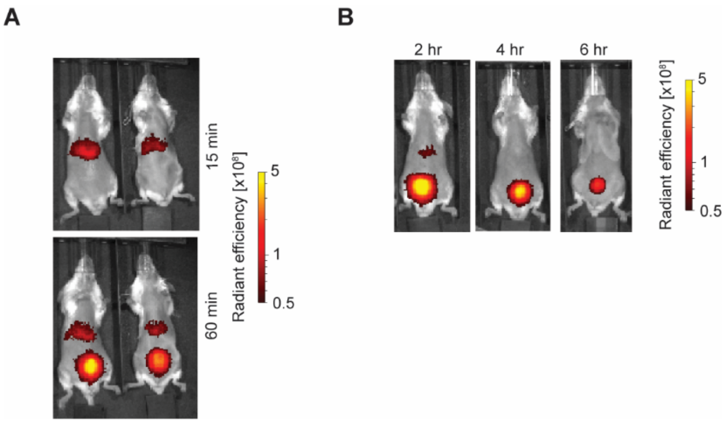


**Figure S4. Supplemental information for in vivo imaging of BALB/c mice after i.v. administration.**

Additional in vivo optical imaging analysis of **PB84-5xAF750. (A)** Replicate fluorescent images of 15- and 60-minute post-injection time points. (B) Representative fluorescent images of 2-, 4-, and 6-hour post-injection time points.

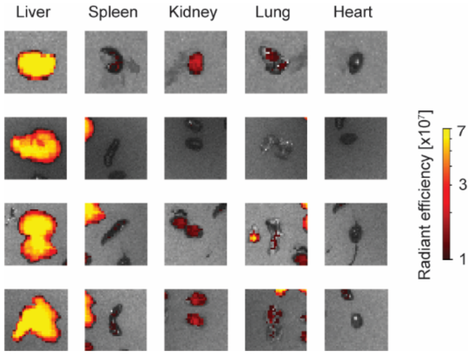


**Figure S5. Supplemental information for ex vivo imaging of BALB/c mice organs after i.v. administration.**

Additional ex vivo optical imaging analysis of **PB84-5xAF750**. Replicate fluorescent images of ex vivo organs harvested 60 minutes post-injection.

### Supplemental Tables

#### Table S1 – Scaffold sequence

| Scaffold Name | Sequence |
|---|---|
| phPB84 | GAGCGCAACGCAATTAATGTGCGCCCTGTAGCGGCGCATTAAGCGCGGCGGGTGTGGTGGTTA<br>CGCGCAGCGTGACCGCTACACTTGCCAGCGCCCTAGCGCCCGCTCCTTTCGCTTTCTTCCCTT<br>CCTTTCTCGCCACGTTCGCCGGCTTTCCCCGTCAAGCTCTAAATCGGGGGCTCCCTTTAGGGTT<br>CCGATTTAGTGCTTTACGGCACCTCGACCCCAAAAAACTTGATTAGGGTGATGGTTCACGTAGT<br>GGGCCATCGCCCTGATAGACGGTTTTTCGCCCTTTGACGTTGGAGTCCACGTTCTTTAATAGTG<br>GACTCTTGTTCCAAACTGGAACAACACTCAACCCTATCTCGGTCTATTCTTTTGATTTATAAGGG<br>ATTTTGCCGATTTCGGCCTATTGGTTAAAAAATGAGCTGATTTAACAAAAATTTAACGCGAATTAC<br>AACCGGGGTACATATGATTGGGGTCTGACGCTCAGTGGAACGAAAACTCACGTTAAGGGATTTT<br>GGTCATGAGATTATCAAAAAGGATCTTCACCTAGATCCTTTTAAATTAAAAATGAAGTTTTAAATC<br>AATCTAAAGTATATATGAGTAAACTTGGTCTGACAGTTACCAATGCTTAATCAGTGAGGCACCTA<br>TCTCAGCGATCTGTCTATTTCGTTCATCCATAGTTGCCTGACTCCCCGTCGTGTAGATAACTACG<br>ATACGGGAGGGCTTACCATCTGGCCCCAGTGCTGCAATGATACCGCGAGACCCACGCTCACCG<br>GCTCCAGATTTATCAGCAATAAACCAGCCAGCCGGAAGGGCCGAGCGCAtAAGTGGTCCTGCAA<br>CTTTATCCGCCTCCATCCAGTCTATTAATTGTTGCCGGGAAGCTAGAGTAAGTAGTTCGCCAGTT<br>AATAGTTTGCGCAACGTTGTTGCCATTGCTACAGGCATCGTGGTGTCACGCTCGTCGTTTGGTA<br>TGGCTTCATTCAGCTCCGGTTCCCAACGATCAAGGCGAGTTACATGATCCCCCATGTTGTGCAA<br>AAAAGCGGTTAGCTCCTTCGGTCCTCCGATCGTTGTCAGAAGTAAGTTGGCCGCAGTGTTATCA<br>CTCATGGTTATGGCAGCACTGCATAATTCTCTTACTGTCATGCCATCCGTAAGATGCTTTTCTGT<br>GACTGGTGAGTACTCAACCAAGTCATTCTGAGAATAGTGTATGCGGCGACCGAGTTGCTCTTGC<br>CCGGCGTCAATACGGGATAATACCGCGCCACATAGCAGAACTTTAAAAGTGCTCATCATTGGAA<br>AACGTTCTTCGGGGCGAAAACTCTCAAGGATCTTACCGCTGTTGAGATCCAGTTCGATGTAACC<br>CACTCGTGCACCCAACTGATCTTCAGCATCTTTTACTTTCACCAGCGTTTCTGGGTGAGCAAAAA<br>CAGGAAGGCAAAATGCCGCAAAAAAGGGAATAAGGGCGACACGGAAATGTTGAATACTCATACT<br>CTTCCTTTTTCAATATTATTGAAGCATTTATCAGGGTTATTGTCTCATGAGCGGATACATATTTGA<br>ATGTATTTAGAAAAATAAACAAATAGGGGTTCCGCGCACATTTCCCCGAAAAGTGCCACCTGAC<br>GTCTAAGAAACCATTATTATCATGACATTAACCTATAAAAATAGGCGTATCACGAGGCCCTTTCG<br>TCGAATTCGTCGTCGTCCCCTCAAACTCTTGGGTGGAGAGGCTATTCGTTTAAGGTCACATCGC<br>ATGTAATTTACTTATTCTCTGTTGTTGAGCCACCCGGGCGCCAGATTTTGTTTAAAGCTTTGTCTC<br>TTAGTTTGTATAGACAGATTCAGAGTGCAAGGTTTCGTTCGCTCGTACCTGGTTTTCCCTGGTTC<br>TTCACAGATAGGATTTGACTTTCTACAACACTTATGCGGCTTCCTACCCGTTTGAAGGCCGATAC<br>AGGTGCTGCGCAAAATGCGGGCGAACATAGAGTATCAAAACAACGCCTTCTAATCTAGGAATAT<br>AGGGAAGATACGTATTTGCTACCATGCTTTCTTGGGTCATTAACGACCAACCTCTTTTCTTTTAAA<br>GTAGGATTGCACAATGAATGAATACACGTGGTCCGATAACTGACCAAGTAACATGGTTATCACTa<br>GATGTCCGCCAGACGTGTGCAAACCAACCCGGGAGTTACGTCACTAATCCTTCGCTACGTCGT<br>GAAGATATTTACTTGTGAATATCGAGGGTAATAAGATAATAGACTGTGACTAGTATTGCCAGACT<br>GTCGCTACCTGCAACACATAACTATCCTGAGGTTACTGCATAGTACTGATTACACCCGAGTCAAA<br>ATTTCTAACTTCTAACATGTACCTAGTAACCAGCTCAATAATTATGTCAGAATATAGCTCTGGGAA<br>CCCTCGGACAATTATGATACACGGTATTAATATCTTGCTTGCGTTAGCCACTTCTCATCTTTGGA<br>TACCGATTCTATTTTGCATAGCAGTTCCTTTTACACATATAAGAATTTCGCCATAGGTATGCTGCA<br>G |

**Table S2 – Staple sequences for PB84**

| Staple Name | Sequence |
|---|---|
| 2 | ATAAAGTTTTGCGTTGCGCTCCTGCAGCATGGATGGAGGCGG |
| 3 | TGCGCTCGCCGCTACAGGGCGCACATTAAGCAGGACCACTTA |
| 4 | GCTGGTTTACACCCGCCGCGCTTAATGCGGCCCTTCCGGCTG |
| 5 | GTCACGCTGCGTTTTTCGTAACCACCATTGCTGATAATTTTTATCTGGAGCCCACGACGGGGATTTTTGTCAGGCAAC |
| 6 | GTGTAGCGAGCCGGCGAACGTGGCGAGAAAGGGCGCTGGCAA |
| 7 | CGGGCGCTAGGAAGGGAAGAAAGCAGCTATAT |
| 8 | TCTGACATGTCCGAGGGTTCCCAGGAAAGGAG |
| 9 | CATTGGTAACTTTTTTGTCAGACCAAGATTTAGAGCTTTTTTTGACGGGGAA |
| 10 | TACTTTAGGGAACCCTAAAGGGAGCCCCCGTTTACTCATATA |
| 11 | TTCATTTTGGTGCCGTAAAGCACTAAATCATTGATTTAAAAC |
| 12 | TCTAGGTGAATCAAGTTTTTTGGGGTCGATAATTTAAAAGGA |
| 13 | GTTGAGTGTTGTTTTTTTCCAGTTTGCACTACGTGAATTTTTCCATCACCCT |
| 14 | CGATGGCCGAACAAGAGTCCACTATTAAACCGTCTATCAGGG |
| 15 | GCGAAAAAGAACGTGGACTCCAACAATCAGTA |
| 16 | CTATGCAGATTTTGACTCGGGTGTGTCAAAGG |
| 17 | GAGATAGGATCAGCTCATTTTTTAACCAAAAAAGAATAGACC |
| 18 | TATAAATCTAGGCCGAAATCGGCATGTTGTAG |
| 19 | AAAGTCAATAGGAAGCCGCATAAGAAATCCCT |
| 20 | AAGATCCTTTTTTTTTTGATAATCTCTTCGCGTTAAATTTTTTTTTTGTTAA |
| 21 | GGTTGTAAATGACCAAAATCCCTTAACGTTCATATGTACCCC |
| 22 | GACCCCAAGAGTTTTCGTTCCACTCCCGTATT |
| 23 | GACGCCGGTGTGGCGCGGTATTATGAGCGTCA |
| 24 | ACTGATTAAGTATGGATGAA |
| 25 | GGTGCCTCCGAAATAGACAGATCGAAAAGCAT |
| 26 | CTTACGGATACTCACCAGTCACAGCTGAGATA |
| 27 | GTTATCTAGGTGAGCGTGGGTCTCGCGGTCTCCCGTATCGTA |
| 28 | GGTAAGCCATCATTGCAGCACTGGAACCGGAG |
| 29 | CTGAATGACGCCTTGATCGTTGGGGGCCAGAT |
| 30 | TCCCGGCAACATTTTTATTAATAGACTACCTATGGCGTTTTTAAATTCTTAT |
| 31 | AATGTGCGGGCGAACTACTTACTCTAGCTCACTTTTCGGGGA |
| 32 | TGTTTATTACGTTGCGCAAACTATTAACTCGGAACCCCTATT |
| 33 | TCAAATATATGCCTGTAGCAATGGCAACATTTCTAAATACAT |
| 34 | AACGACGAGCGTTTTTTGACACCACGGTATCCGCTCATTTTTTGAGACAATA |
| 35 | AGCCATACCATCATGTAACT |
| 36 | CATTTCCGTGTTTTTTCGCCCTTATTTTTTTTGCACATTTTTACATGGGGGA |
| 37 | CATTTTGCAGGACCGAAGGAGCTAACCGCCCCTTTTTTGCGG |

| | |
|---|---|
| 38 | CTCACCCACTTACTTCTGACAACGATCGGCTTCCTGTTTTTG |
| 39 | AAGTAAAAGAGTGATAACACTGCGGCCAAGAAACGCTGGTGA |
| 40 | CTCGGTCGCCGTTTTTCATACACTATATGCAGTGCTGTTTTTCCATAACCAT |
| 41 | AGAGAATTTCTCAGAATGACTTGGTTGAGTGGCATGACAGTA |
| 42 | GCAAGAGCAAAGTTCTGCTA |
| 43 | GATGCTGAAGATTTTTTCAGTTGGGTTCCAATGATGATTTTTGCACTTTTAA |
| 44 | GAACGTTTGCACGAGTGGGTTACATCGAATTTTCGCCCCGAA |
| 45 | CTTGAGAGCTGGATCTCAACAGCGTGAATCTG |
| 46 | TCTATACAACGAAACCTTGCACTCGTAAGATC |
| 47 | GAGTATTCAAACCCTGATAA |
| 48 | AAGAGTATATGCTTCAATAATATTACATGCGA |
| 49 | TGTGACCTAGAGAATAAGTAAATTGAAAAAGG |
| 50 | TTATCGGACCATTTTTCGTGTATTCAGGTTTCTTAGATTTTTCGTCAGGTGG |
| 51 | CCTACTTTGGTTAATGTCATGATAATAATTTCATTGTGCAAT |
| 52 | TTGGTCGTCGTGATACGCCTATTTTTATAAAAAGAAAAGAGG |
| 53 | AAGCATGGACGAATTCGACGAAAGGGCCTTAATGACCCAAGA |
| 54 | CAAAGCTTTAATTTTTACAAAATCTGCAAGAGTTTGATTTTTGGGGACGACGTAGCAAATACGTTTT TTATCTTCCCT |
| 55 | TCTCCACCGCGCCCGGGTGGCTCAACAACTAAACGAATAGCC |
| 56 | AACTAAGAGAGTACGAGCGA |
| 57 | TGCGCAGCACCTTTTTTGTATCGGCCTGAAGAACCAGTTTTTGGAAAACCAG |
| 58 | ATCCTATCTGTTCAAACGGG |
| 59 | GCCCGCATTTATATTCCTAG |
| 60 | CTATGTTCATTAGAAGGCGTTGTTATTACCCT |
| 61 | CGATATTCCACAGTCTATTATCTTTTGATACT |
| 62 | ACTGCTATATAACCATGTTACTTGGTCAGATGTGTAAAAGGA |
| 63 | GGTATCCAACGTCTGGCGGACATCTAGTGGCAAAATAGAATC |
| 64 | GCTAACGCAACTCCCGGGTTGGTTTGCACAAGATGAGAAGTG |
| 65 | GTAGCGAAGGATTTTTTTAGTGACGTAAGCAAGATATTTTTTTAATACCGTG |
| 66 | TTCACGACCGACAGTCTGGCAATACTAGTACAAGTAAATATC |
| 67 | CTGGTTACTAGTTTTTGTACATGTTAATAGTTATGTGTTTTTTTGCAGGTAG |
| 68 | TAACCTCAGGGAAGTTAGAA |
| 69 | AATTATTGAGTATCATAATT |

**Table S3 – Staple sequences for PB84-5xAF750**

| Staple Name | Sequence |
|---|---|
| 2 | ATAAAGTTTTGCGTTGCGCTCCTGCAGCATGGATGGAGGCGG |
| 3 | TGCGCTCGCCGCTACAGGGCGCACATTAAGCAGGACCACTTA |
| 4 | GCTGGTTTACACCCGCCGCGCTTAATGCGGCCCTTCCGGCTG |
| 5 | GTCACGCTGCGTTTTTCGTAACCACCATTGCTGATAATTTTTATCTGGAGCCCACGACGGGGATTTTTGTCAGGCAAC |
| 6 | GTGTAGCGAGCCGGCGAACGTGGCGAGAAAGGGCGCTGGCAA |
| 7 | CGGGCGCTAGGAAGGGAAGAAAGCAGCTATAT |
| 8 | TCTGACATGTCCGAGGGTTCCCAGGAAAGGAG |
| 9 | CATTGGTAACTTTTTTGTCAGACCAAGATTTAGAGCTTTTTTTGACGGGGAA |
| 10 | TACTTTAGGGAACCCTAAAGGGAGCCCCCGTTTACTCATATA |
| 11 | TTCATTTTGGTGCCGTAAAGCACTAAATCATTGATTTAAAAC |
| 12 | TCTAGGTGAATCAAGTTTTTTGGGGTCGATAATTTAAAAGGA |
| 13 | GTTGAGTGTTGTTTTTTTCCAGTTTGCACTACGTGAATTTTTCCATCACCCT |
| 14-AF750 | AF750TTCGATGGCCGAACAAGAGTCCACTATTAAACCGTCTATCAGGG |
| 15 | GCGAAAAAGAACGTGGACTCCAACAATCAGTA |
| 16 | CTATGCAGATTTTGACTCGGGTGTGTCAAAGG |
| 17 | GAGATAGGATCAGCTCATTTTTTAACCAAAAAAGAATAGACC |
| 18 | TATAAATCTAGGCCGAAATCGGCATGTTGTAG |
| 19 | AAAGTCAATAGGAAGCCGCATAAGAAATCCCT |
| 20 | AAGATCCTTTTTTTTTTGATAATCTCTTCGCGTTAAATTTTTTTTTTGTTAA |
| 21-AF750 | AF750TTGGTTGTAAATGACCAAAATCCCTTAACGTTCATATGTACCCC |
| 22 | GACCCCAAGAGTTTTCGTTCCACTCCCGTATT |
| 23 | GACGCCGGTGTGGCGCGGTATTATGAGCGTCA |
| 24 | ACTGATTAAGTATGGATGAA |
| 25 | GGTGCCTCCGAAATAGACAGATCGAAAAGCAT |
| 26 | CTTACGGATACTCACCAGTCACAGCTGAGATA |
| 27 | GTTATCTAGGTGAGCGTGGGTCTCGCGGTCTCCCGTATCGTA |
| 28 | GGTAAGCCATCATTGCAGCACTGGAACCGGAG |
| 29 | CTGAATGACGCCTTGATCGTTGGGGGCCAGAT |
| 30 | TCCCGGCAACATTTTTATTAATAGACTACCTATGGCGTTTTTAAATTCTTAT |
| 31-AF750 | AF750TTAATGTGCGGGCGAACTACTTACTCTAGCTCACTTTTCGGGGA |
| 32 | TGTTTATTACGTTGCGCAAACTATTAACTCGGAACCCCTATT |
| 33 | TCAAATATATGCCTGTAGCAATGGCAACATTTCTAAATACAT |
| 34 | AACGACGAGCGTTTTTTGACACCACGGTATCCGCTCATTTTTTGAGACAATA |
| 35 | AGCCATACCATCATGTAACT |
| 36 | CATTTCCGTGTTTTTTCGCCCTTATTTTTTTTGCACATTTTTACATGGGGGA |
| 37-AF750 | AF750TTCATTTTGCAGGACCGAAGGAGCTAACCGCCCCTTTTTTGCGG |

| | |
|---|---|
| 38 | CTCACCCACTTACTTCTGACAACGATCGGCTTCCTGTTTTTG |
| 39 | AAGTAAAAGAGTGATAACACTGCGGCCAAGAAACGCTGGTGA |
| 40 | CTCGGTCGCCGTTTTTCATACACTATATGCAGTGCTGTTTTTCCATAACCAT |
| 41 | AGAGAATTTCTCAGAATGACTTGGTTGAGTGGCATGACAGTA |
| 42 | GCAAGAGCAAAGTTCTGCTA |
| 43 | GATGCTGAAGATTTTTTCAGTTGGGTTCCAATGATGATTTTTGCACTTTTAA |
| 44 | GAACGTTTGCACGAGTGGGTTACATCGAATTTTCGCCCCGAA |
| 45 | CTTGAGAGCTGGATCTCAACAGCGTGAATCTG |
| 46 | TCTATACAACGAAACCTTGCACTCGTAAGATC |
| 47 | GAGTATTCAAACCCTGATAA |
| 48 | AAGAGTATATGCTTCAATAATATTACATGCGA |
| 49 | TGTGACCTAGAGAATAAGTAAATTGAAAAAGG |
| 50 | TTATCGGACCATTTTTCGTGTATTCAGGTTTCTTAGATTTTTCGTCAGGTGG |
| 51 | CCTACTTTGGTTAATGTCATGATAATAATTTCATTGTGCAAT |
| 52 | TTGGTCGTCGTGATACGCCTATTTTTATAAAAAGAAAAGAGG |
| 53 | AAGCATGGACGAATTCGACGAAAGGGCCTTAATGACCCAAGA |
| 54 | CAAAGCTTTAATTTTTACAAAATCTGCAAGAGTTTGATTTTTGGGGACGACGTAGCAAATACGTTT TTTATCTTCCCT |
| 55 | TCTCCACCGCGCCCGGGTGGCTCAACAACTAAACGAATAGCC |
| 56 | AACTAAGAGAGTACGAGCGA |
| 57 | TGCGCAGCACCTTTTTTGTATCGGCCTGAAGAACCAGTTTTTGGAAAACCAG |
| 58 | ATCCTATCTGTTCAAACGGG |
| 59 | GCCCGCATTTATATTCCTAG |
| 60 | CTATGTTCATTAGAAGGCGTTGTTATTACCCT |
| 61 | CGATATTCCACAGTCTATTATCTTTTGATACT |
| 62 | ACTGCTATATAACCATGTTACTTGGTCAGATGTGTAAAAGGA |
| 63 | GGTATCCAACGTCTGGCGGACATCTAGTGGCAAAATAGAATC |
| 64-AF750 | AF750TTGCTAACGCAACTCCCGGGTTGGTTTGCACAAGATGAGAAGTG |
| 65 | GTAGCGAAGGATTTTTTTAGTGACGTAAGCAAGATATTTTTTTAATACCGTG |
| 66 | TTCACGACCGACAGTCTGGCAATACTAGTACAAGTAAATATC |
| 67 | CTGGTTACTAGTTTTTGTACATGTTAATAGTTATGTGTTTTTTTGCAGGTAG |
| 68 | TAACCTCAGGGAAGTTAGAA |
| 69 | AATTATTGAGTATCATAATT |

**Table S4 – Supplemental information for cytokine induction in BALB/c mice after i.v. administration: averages**

| Averages | | mIFNa (pg/ml) | mIFNb (pg/ml) | mIFNy (pg/ml) | mIL-1b (pg/ml) | mIL-6 (pg/ml) | mIL-12 (pg/ml) | mTNFa (pg/ml) | mCXCL1 (pg/ml) | mCXCL2 (pg/ml) |
|---|---|---|---|---|---|---|---|---|---|---|
| 3 h | PBS | 62.48 | 12.44 | 33.96 | 0.20* | 0.20* | 47.04 | 0.10* | 22.74 | 38.42 |
| | PB84 | 57.87 | 10.55 | 23.71 | 0.20* | 9.17 | 23.48 | 0.10* | 20.53 | 163.50 |
| | LPS | 20.59 | 10.59 | 14.86 | 70.71 | 26152.60 | 146.29 | 21.60 | 48893.87 | 841.79 |
| 24 h | PBS | 42.28 | 10.68 | 15.30 | 0.20* | 0.20* | 7.02 | 0.10* | 7.32 | 112.20 |
| | PB84 | 52.20 | 8.67 | 0.20* | 0.27 | 0.20* | 121.47 | 3.09 | 9.50 | 28.70 |
| | LPS | 87.81 | 14.30 | 56.69 | 5.21 | 67.85 | 47.39 | 3.02 | 525.25 | 139.14 |

* All replicates below the lower limit of detection – lower limit of detection reported.

**Table S5 – Supplemental information for cytokine induction in BALB/c mice after i.v. administration: relative standard error of the mean**

| Relative SEM | | mIFNa (pg/ml) | mIFNb (pg/ml) | mIFNy (pg/ml) | mIL-1b (pg/ml) | mIL-6 (pg/ml) | mIL-12 (pg/ml) | mTNFa (pg/ml) | mCXCL1 (pg/ml) | mCXCL2 (pg/ml) |
|---|---|---|---|---|---|---|---|---|---|---|
| 3 h | PBS | 0.06 | 0.19 | 0.65 | n.d.* | n.d.* | 0.94 | n.d.* | 0.45 | 0.67 |
| | PB84 | 0.26 | 0.04 | 0.89 | n.d.* | 0.98 | 0.76 | n.d.* | 0.21 | 0.59 |
| | LPS | 0.75 | 0.05 | 0.72 | 0.50 | 0.32 | 0.24 | 0.28 | 0.25 | 0.26 |
| 24 h | PBS | 0.88 | 0.02 | 0.50 | n.d.* | n.d.* | 0.62 | n.d.* | 0.53 | 0.47 |
| | PB84 | 0.51 | 0.14 | n.d.* | 0.27 | n.d.* | 0.83 | 0.85 | 0.59 | 0.70 |
| | LPS | 0.26 | 0.14 | 0.70 | 0.96 | 0.88 | 0.71 | 0.71 | 0.78 | 0.84 |

* All replicates below the lower limit of detection – lower limit of detection reported.
